# Concise functional enrichment of ranked gene lists

**DOI:** 10.1101/2023.06.30.547164

**Authors:** Xinglin Jia, An Phan, Claus Kadelka

## Abstract

Genome-wide expression data has become ubiquitous within the last two decades. Given such data, functional enrichment methods identify functional categories (e.g., biological processes) that preferentially annotate differentially expressed genes. However, many existing methods operate in a binary manner, disregarding valuable information contained in the gene ranking. The few methods that consider the ranking often return redundant or non-specific functional categories.

To address these limitations, we developed a novel method called Concise Ranked Functional Enrichment (CRFE), which effectively leverages the ranking information in gene expression data to compute a non-redundant set of specific functional categories that are notably enriched for highly ranked genes. A particularly useful feature of CRFE is a tunable parameter that defines how much focus should be given to the most highly ranked genes. Using four treatment-control RNA-seq datasets, we compared the performance of CRFE with the two most widely used types of functional enrichment methods, Gene Set Enrichment Analysis and over-representation analysis. We evaluated the methods based on their ability to utilize ranking information, generate non-redundant results, and return functional categories with high information content. CRFE excelled in all evaluated criteria, outperforming the existing methods, each of which exhibits deficiencies in at least one aspect. Using lung adenocarcinoma data, we further showed that the functional categories identified by CRFE are biologically meaningful.

In conclusion, CRFE computes an informative set of functional categories that summarizes genome-wide expression data. With its superior performance over existing methods, CRFE harbors great promise to become a widely used functional enrichment method.

**Author summary:** Given a list of differentially expressed genes as input, functional enrichment methods reveal which functional categories (e.g., biological processes) were likely activated by the cell and are responsible for the differential expression. We developed a new such method, called Concise Ranked Functional Enrichment (CRFE), which addresses the limitations of current approaches by incorporating gene ranking information to compute a concise and specific set of enriched functional categories. Using four treatment-control RNA-seq datasets, we evaluate how well CRFE and the two currently most widely used methods perform in three criteria. We find that CRFE outperforms each of the alternative methods in at least one of the evaluated criteria, demonstrating its superiority. A high-level interpretation of the functional categories identified by CRFE for lung adenocarcinoma datasets highlights its usefulness for experimentalists. Overall, CRFE harnesses the power of ranked gene lists to generate a focused and non-redundant set of enriched functional categories. Our study positions CRFE as a promising method for functional enrichment analysis, with the potential to advance research in this field.

## Introduction

The advent of next-generation sequencing technology has ushered in a deluge of high-throughput data in biological studies [1, 2]. Over the past two decades, scientists have become able to conduct comprehensive profiling studies on the genome, epigenome, transcriptome, metabolome, proteome, etc, at increasingly affordable costs [3, 4]. These omics studies typically generate extensive lists of biological components, such as genes, proteins, and metabolites, which can be ranked based on their abundance, fold change, or other relevant attribute. The interpretation of such data presents a fundamental and challenging bioinformatics task [5–7]. Functional enrichment methods are usually employed to make sense of these long lists. In the most common use case, these methods are applied to long lists of genes, ranked by differential expression values from treatment-control transcriptome study (e.g., RNA-seq). The output typically consists of a selection of functional categories that annotate a surprisingly large number of highly ranked genes [8, 9]. The returned functional categories provide a biological explanation of the gene expression changes, facilitating a more comprehensive understanding of the underlying biology. In this study, we adhere to this most common use case. Functional enrichment methods are, however, versatile and applicable to other omics data [7, 10–12]. Functional enrichment methods require an annotation database such as the Gene Ontology (GO), which describes which genes are involved in which functional categories. The GO is a controlled vocabulary of cellular components, molecular functions, and biological processes, with a directed acyclic graph structure [13]. Within this graph, parent nodes correspond to broader functional categories, while child nodes represent more specific functional categories. Each functional category within the GO is associated with a set of genes. Parent nodes encompass all genes from their respective child nodes. Consequently, child functional categories are generally more informative (i.e., possess a higher information content) than their respective parent categories [14–16].

Recent citation counts indicate most researchers rely on two families of functional enrichment methods: over-representation analyses (ORA) and derivatives of the gene set enrichment analysis (GSEA) method [6, 17, 18]. Part of their popularity likely stems from their streamlined implementation through ready-to-use software packages and/or R libraries. [18–22]. However, all popular methods suffer from at least one shortcoming. ORA-based methods assume gene independence, implying that perturbation of one gene does not correlate with perturbation of any other gene. Since genes are commonly co-expressed, this simplifying assumption is incorrect. Additionally, all ORA-based methods operate in a binary mode; they classify genes as perturbed and unperturbed based on a user-defined cutoff, and identify functional categories that annotate many perturbed and few unperturbed genes. This approach disregards valuable information contained in the gene ranking. By using the full information provided by the ranking, it should be possible to identify a set of functional categories that explains the highly ranked genes particularly well, thereby maximizing the potential insights gained from the analysis.

The other popular method, GSEA and its derivatives, operates on ranked gene lists [18, 23–26]. These methods consist of four major steps: calculating local (gene-level) and global (category-level) statistics, determining significance, and adjusting for multiple testing [18]. Users have the freedom to choose methods for each step based on their data type and prior analysis, but this flexibility can lead to inconsistent outputs, causing different users starting with the same expression data to make entirely different discoveries. Although the derivatives exist to address specific GSEA issues, the lack of a comprehensive benchmark and a universally recognized standard for their usage remains unresolved [5, 27].

Furthermore, GSEA-based methods, along with ORA or any other methods that regard each functional category individually require a multiple testing correction. This correction can lead to a limited number of functional categories that are considered significantly enriched [26]. The assumption of gene and gene set independence also fails to consider the overlap among categories that arises from the hierarchical structure of annotation databases. As a result, the list of returned functional categories may be highly redundant, in the sense that the sets of genes annotated to the returned categories are similar. Manually selecting a non-redundant and meaningful subset of functional categories can be a challenging task. To aid in this interpretation, methods such as REVIGO have been developed to summarize lengthy lists of functional categories [24, 28, 29]. REVIGO works by clustering highly similar GO categories based on semantic similarity measures; the categories that remain in the list are cluster representatives. These methods can be applied post-hoc to the output of functional enrichment methods to avoid redundant results.

This review of the existing literature highlights that an ideal functional enrichment method for ranked gene lists should fulfill three key criteria. Firstly, the method should be able to identify functional categories that annotate a substantial number of differentially expressed genes, with a preference for highly ranked genes. This ensures that the method captures the most relevant biological insights by focusing on genes that exhibit significant changes in expression. Secondly, the method should provide a non-redundant set of categories, meaning that the identified categories should share hardly any gene annotations. This avoids repetitive information and allows for a more concise and focused interpretation of the results. Lastly, it is advantageous for the method to prioritize specific categories over those with a large number of gene annotations. Categories with fewer annotations tend to be more precisely defined and offer more informative insights into the underlying biological processes. This preference for specificity ensures that the method highlights well-defined biological processes that are directly associated with the differentially expressed genes.

Model-based Gene Set Analysis (MGSA) is one of a few functional enrichment methods that consider sets of functional categories, rather than each category individually [30]. By finding the set of functional categories that collectively best explains a given expression dataset, MGSA manages to return a smaller non-redundant set of functional categories. MGSA operates, however, only in binary mode and thus fails to leverage the valuable information provided by gene ranking. To overcome this limitation, we propose a novel method called Concise Ranked Functional Enrichment (CRFE). As MGSA, CRFE classifies the genes into perturbed and unperturbed based on a user-defined threshold. Contrary to MGSA, CRFE considers the specific ranking of all perturbed genes. It returns a concise and non-redundant set of functional categories, which prioritizes the inclusion of highly ranked genes among the returned categories.

In this manuscript, we introduce CRFE, and using four treatment-control RNA-seq datasets and three functional enrichment performance metrics, we assess the effectiveness of CRFE compared to the two widely used methods, ORA and GSEA, as well as MGSA, which CRFE generalizes. We show that CRFE performs at least as well as each other method in all three metrics, and strictly better than each other method in at least one metric. We conclude with a brief biological interpretation of the top functional categories returned by CRFE for two human lung adenocarcinoma RNA-seq datasets, demonstrating the practical utility and effectiveness of CRFE in deciphering the biological implications of gene expression data.

## Methods

### Generative model

We employ a generative model, as in MGSA [30], that explains how biological experiments yield lists of thousands of genes, proteins, etc, ranked by some measured value. For ease of exposition, we describe the model in the context of a treatment-control RNA-seq experiment, which is the most frequent use case of functional enrichment methods. Here, the genes are ranked by differential expression value (e.g., fold change) between treatment and control. We model the response of genes in the experiment as the result of the activation of several functional categories due to the treatment.

We assume that the experiment seeks to detect gene states based on differential expression, and that a gene can be in one of two general states, “perturbed,” or “unperturbed”. The true state of every gene is hidden. We model the results of the experiment by assuming that, in the absence of any noise, every gene that is in the “perturbed” state is annotated to at least one activated functional category. However, since experimental data contains noise, we further assume that unknown false positive and false negative rates determine the observed state of every gene.

We now describe the generative model formally (see Figure 1 for an illustration). We use biological processes in the Gene Ontology to represent functional categories. Let *C* denote the set of all biological processes. In response to the treatment, a cell activates an unknown proportion *q* ∈ (0, 1) of these processes independently. We denote the set of all activated processes by *C* ⊂ *C*.

**Fig 1.**
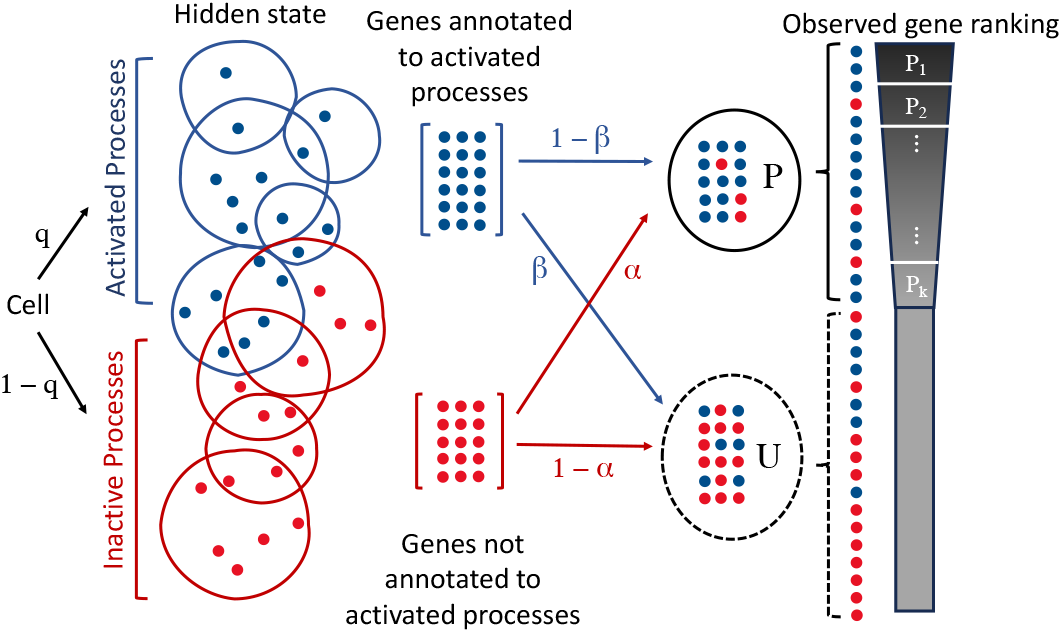
Illustration of the generative model. In response to the treatment, the cell activates a proportion *q* of biological processes. Genes annotated to the activated processes are observed perturbed or unperturbed in a differential expression analysis, with true positive rate 1 *− β* and false negative rate *β*, respectively. Likewise, genes not annotated to any activated processes are observed perturbed or unperturbed with false positive rate *α*, and true negative rate 1 *− α*, respectively. For this specific illustration, we use the values *α* = 1*/*5, *β* = 1*/*3, and *q* = 5*/*9.

Genes annotated to the activated processes exhibit varying degrees of expression: some genes are highly expressed due to the treatment, and are therefore more likely to exhibit a high differential expression value. These genes are observed perturbed; other annotated genes may display no difference in expression levels and are observed unperturbed. This difference in expression gives rise to a stratification of the genes into two sets: *P*, the perturbed genes, and *U*, the unperturbed genes. We assume that genes annotated to an activated process are observed unperturbed with probability *β*, the false negative rate. Similarly due to random noise, we assume that all genes not annotated to any activated process are observed perturbed with probability *α*, the false positive rate. We further assume that *α* and *β* are identical and independent for all genes.

This generative model can also be represented as a Bayesian network, wherein the sets *C* and *C − C* constitute a hidden layer, and the gene sets *P* and *U* form the observed layer. The hidden layer is conditioned on the parameter *q*, while the observed layer is conditioned on the hidden layer as well as the parameters *α* and *β*.

### Algorithm

CRFE aims to recover the set of activated functional categories *C* that gave rise to the observed ranked gene list, which can be divided into perturbed and unperturbed genes based on a user-defined ranking cutoff *τ*. Common cutoffs include a 2-fold or 1.5-fold change in expression values. In an ideal scenario, CRFE finds a set of categories *C* such that all perturbed genes would be annotated to at least one category *c* ∈ *C*, while none of the unperturbed genes would be annotated to any category *c* ∈ *C*. However due to noise, this ideal solution is typically unachievable. We thus need to find a *C* that best explains the observed gene list and define what “best” means.

For a given *C*, let *E*(*C*) be the set of genes annotated to at least one category *c ∈ C*. We call elements in *E*(*C*) explained genes, and say that *C* explains the set of genes *E*(*C*). Further, let *N* = *N* (*C*) contain all unexplained genes, i.e., those not annotated to any category *c ∈ C*.

Higher-ranked perturbed genes are more likely to represent true perturbations than genes just above the ranking cutoff *τ*. It is therefore more important to explain highly ranked genes. To achieve this emphasis, we divided the set *P* of perturbed genes into *k* equally-sized subsets, *P*_1_, …, *P*_*k*_, based on the ranking and introduce subset-specific weights *w*_*i*_ later. Genes in *P*_*i*_ are higher ranked (i.e., more differentially expressed) than those in *P*_*i*+1_. For computational efficiency and to prevent the loss of even marginal information provided by the ranking, we considered *k* = |*P* |, i.e., each perturbed gene forms its own subset. However, in theory, any value *k ∈* 1, 2, …, |*P* | is feasible, with *k* = 1 corresponding to the MGSA approach.

We can define four disjoint types of gene sets:

i. *EP*_*i*_ = *E*(*C*) *∩ P*_*i*_ contains the perturbed genes in subset *P*_*i*_ that are annotated to at least one category *c ∈ C*,
ii. *NP*_*i*_ = *N* (*C*) *∩ P*_*i*_ contains the perturbed genes in subset *P*_*i*_ that are not annotated to any category *c ∈ C*,
iii. *EU* = *E*(*C*) *∩ U* contains the unperturbed genes that are annotated to at least one category *c ∈ C*, and
iv. *NU* = *N* (*C*) *∩ U* contains the unperturbed genes that are not annotated to any category *c ∈ C*.

Each of these gene sets is a function of *C*. To simplify notation, we will omit the dependence on *C*.

The joint probability of observing a ranked list of genes *P* = (*P*_1_, …, *P*_*k*_), *U* given a set *C* of activated categories, as well as parameters *α, β*, and *q* is then

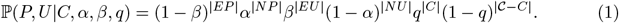

Since 0 *< q <<* 1, the parameter *q* penalizes increases in the size of *C*. This joint probability is also the likelihood of (*C, α, β, q*) given the ranked gene list, denoted as *L*(*C, α, β, q*|*P, U*). MGSA maximizes this likelihood function. With *P* split into *P*_*i*_, *i* = 1, …, *k*, we can rewrite Eq 1 as

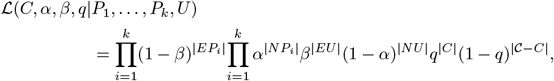

which can be modified by including subset-specific weights, enabling CRFE to put more emphasis on explaining highly-perturbed genes.

### Weights

A perturbed gene contributes 1 *− β* to the likelihood function (Eq 1) if it is explained and *α* if it is not. If *α, β ≤* 0.5, then 1 *− β ≥ α*, implying that it is always beneficial (i.e., *L* increases) to find a *C* that explains perturbed genes. To increase the importance of explaining highly ranked perturbed genes, we introduced weights to replace 1 *− β* by 1 *− βw*_*i*_ and *α* by *αw*_*i*_ for genes in *P*_*i*_. Requiring *w*_*i*_ *< w*_*i*+1_ ensures it is more important to explain a higher-ranked gene. The likelihood function with these weights is

*L*(*C, α, β, q*|*P*_1_, …, *P*_*k*_, *U*)

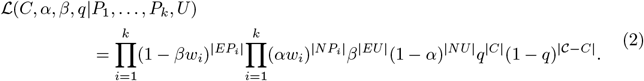

We designed the weights *w*_*i*_ with specific properties to facilitate the interpretation and analysis (Figure 2).

**Fig 2.**
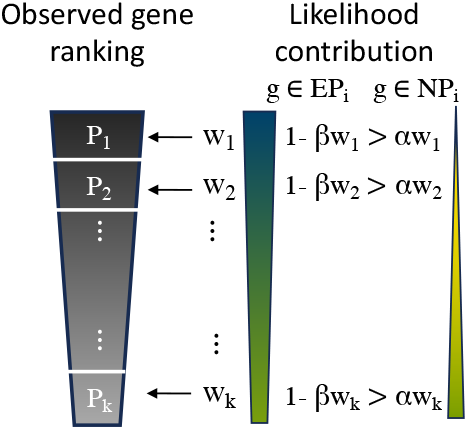
Weights in CRFE. To prioritize the explanation of highly perturbed genes by CRFE, we designed the weights such that the likelihood function is “rewarded more” if a highly perturbed gene is annotated to at least one functional category in the explanatory set *C* than if a less perturbed gene is explained by *C*. Note that even for the least perturbed gene (in *P*_*k*_) the likelihood is higher if it is explained (i.e., if it is in *EP*_*k*_). This holds as long as 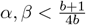.

1. We assumed the weights increase at a fixed linear rate. That is,

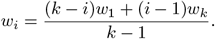
2. We introduced a hyper-parameter *b*, which describes the difference between the highest and lowest weight,

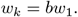

If *b* = 1, all perturbed subsets are assigned equal weights, which is equivalent to the approach used in MGSA. If *b >* 1, the likelihood decreases more if a highly ranked perturbed gene is not explained. We call *b* the belief parameter, as the choice of a higher value for *b* intuitively translates to an experimentalist having greater confidence (“belief”) that the genes measured to be highly differentially expressed are the most relevant to the experiment.
3. We centered the weights at 1 to achieve *w*_*i*_ *∈* (0, 2) and ultimately, 1 *− βw*_*i*_, *αw*_*i*_ *∈* (0, 1). That is,

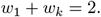

These conditions lead to a linear system with a unique solution (*w*_1_, …, *w*_*k*_) given by

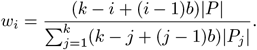

If each perturbed gene forms its own subset (i.e., *k* = |*P* |, |*P*_*i*_| = 1), as used in our implementation of CRFE, this simplifies to

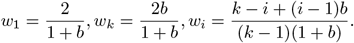

When learning the parameters in the Markov chain Monte Carlo (MCMC) process (defined below), the possible range for *α* and *β* needs to be restricted in order to reward CRFE for explaining any perturbed gene. That implies *αw*_*i*_ *≤* 1 *− βw*_*i*_ for all *i* = 1, …, *k*. Since *w*_*i*_ is increasing in *i*, this reduces to *αw*_*k*_ *≤* 1 *− βw*_*k*_. In the MCMC process, *α, β* are independently updated within the range [0, *r*_max_]. We thus require

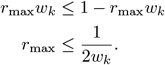

If *k* = |*P* |, this yields 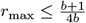, which approaches 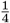 for very large choices of the belief parameter *b*.

### Markov chain Monte Carlo Optimization

Let *θ* = (*C, α, β, q*) and let Θ = {0, 1}^*C*^ *× 𝒜 × 𝒞 × 𝒬* be the solution space of *θ* where *𝒜, 𝒞, 𝒬* contain all possible values of *α, β, q*, respectively. Finding a set *C*^*\**^ of functional categories that “best” explains the data reduces to maximizing the likelihood function (Eq 2). That is,

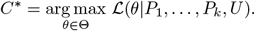

Even for a fixed choice of *α, β, q*, the solution space Θ has a size of 2^|*C*|^, which is too large to be explored exhaustively. Therefore, we employed a Markov Chain Monte Carlo (MCMC) method to estimate *C*^*\**^ and the parameters (*α, β, q*). For convenience, we maximized the log-likelihood function,

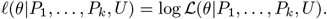

We initialized the set of activated processes as the empty set, *C*_0_ = *ø*. We further set *α* = 0.1, *β* = 0.25, *q* = 1*/*|*C*| (since we learn *α, β*, the initial choices do not matter). During each iteration *j ≥* 1, we generated a new proposal set 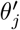 by one of the following three actions:

1. with a probability of 80%, we updated *C*. That is,
  a. we added an unselected functional category (from *C − C*_*j−*1_) to the current set *C*_*j−*1_, or
  b. we deleted a category from the current set *C*_*j−*1_, or
  c. we swapped a category in *C*_*j−*1_ with an unselected category.
2. with a probability of 10%, we updated *α*.
3. with a probability of 10%, we updated *β*.

As we ran the Markov chain for many iterations (i.e., until sufficient convergence had occurred), the specific probabilities assigned to each action did not matter.

There are a total of |*C*| + |*C*_*j−*1_||*C − C*_*j−*1_| possible proposals for the first action. We chose each proposal with equal probability. For computational efficiency and to obtain a finite-state Markov chain, we considered only 20 choices for *α* and *β*.We set

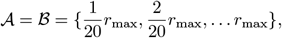

and picked any value with equal probability. We updated *q* whenever a functional category was added or deleted from the current set *C*_*j−*1_ and used an upper bound of min(|*C*_*j−*1_|, 20)*/*|*C*|, similar to the approach used in MGSA. This upper limit for *q* ensured that the penalty for adding categories remained sufficiently high.

We accepted the proposed change with probability

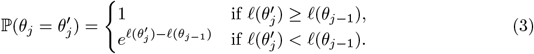

Otherwise, the proposal was rejected and *θ*_*j*_ remained unchanged. In other words, if the likelihood of the proposed set was higher than the likelihood of the current set, the proposal was always accepted. However, if the likelihood of the proposed set was lower, we still accepted the proposal with a positive probability, which is inversely proportional to the difference in likelihoods. This allowed the search process to occasionally explore regions with lower likelihoods, enabling the escape from local maxima. The Markov chain generated by this process is finite, irreducible, and aperiodic. Each state can be reached with a positive probability from any other state within |*C*| + 2 steps. These properties ensure convergence to a stationary distribution that can be sampled from, providing us with the inferred set of activated functional categories.

To allow the Markov chain to reach a stationary distribution, we incorporated a burn-in period of *m* = 50, 000 iterations, after which we recorded the current set *θ*_*j*_ at each iteration for an additional *n* = 50, 000 iterations. To determine the posterior probability of a functional category *c ∈ C*, we calculated the fraction of recorded steps in which the current set *C*_*j*_ contained *c*. Mathematically, the posterior probability of *c ∈ C* is defined as

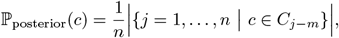

where *C*_*j−m*_ represents the *j*th set recorded after the burn-in period. We ranked all functional categories by their posterior probability, which indicates the importance of each category in explaining the observed data, and returned all categories with a positive posterior probability.

## Data preprocessing

### Gene Ontology annotations

We used the human Gene Ontology annotation file (GAF), retrieved from the Gene Ontology (GO) Consortium in December 2022 [31], which records all leaf annotations in the biological process aspect of human genes. The annotations were then propagated using the Gene Ontology structure stored in an OBO file (containing the relationships between GO categories) to respect the true path rule, i.e., any gene annotated to a GO category is also annotated to the ancestors of that GO category. Next, we combined all biological processes which annotate exactly the same genes (e.g., “growth” and “developmental growth” are combined into “ growth & developmental growth”) while other processes were retained. We did this to improve the performance and consistency of the MCMC process. This way, the Markov chain is not forced to choose one of possibly many equal categories arbitrarily (and assign potentially a high posterior probability) and discard the others (and assign a low posterior probability).

Consequently, an annotation file, including combined GO categories and genes explained by each corresponding category, is used as inputs to the functional enrichment methods.

### Datasets

We compared the performance of different functional enrichment methods using four published RNA-seq treatment-control datasets: two lung adenocarcinoma datasets (GSE40419, GSE87340, from the Gene Expression Omnibus (GEO) database) and two COVID-19-related datasets sampled from the lung and colon of COVID patients and healthy individuals (E-ENAD-46, from the ExpressionAtlas database) [32–34]. The first dataset (GSE40419) comes from an RNA-seq experiment identifying somatic point mutations and transcriptional variants, containing 87 cancer samples and 77 adjacent normal lung tissue samples from primary lung adenocarcinoma patients. Expression levels of more than 17,000 genes are publicly available as Reads Per Kilobase of transcript per Million mapped reads (RPKM). The second cancer dataset (GSE87340) is obtained from an RNA-seq experiment performed on 27 lung adenocarcinomas to examine the recurrent gene mutations and correlation with pathway deregulation and patient outcome. Paired samples were taken from the primary tumor and the unaffected normal lung tissue from stage I adenocarcinoma never-smokers. The raw counts of more than 21,000 genes are available at the GEO database. The third dataset (E-ENAD-46) contains samples taken from colon tissue (denoted EENADCL from now on) and lung tissue (denoted EENADLU from now on) of nine COVID-19 patients. The control samples (not COVID-19 infected) were obtained from healthy colon and lung tissues of cancer patients.

For the first two datasets, we performed differential gene expression analysis using DESeq2 [35] to get a list of genes ranked by *log*2-fold change (using normal samples as the baseline). For the two COVID-19 datasets, we obtained the published *log*2-fold change data and corresponding *p*-values from ExpressionAtlas [36]. The *log*2-fold change values were rounded to one decimal place only and therefore exhibited many ties, leading to arbitrariness in the gene order and negative effects on the outcomes of functional enrichment methods. To increase the signal in the data, we broke most ties by ranking by *p*-values as a secondary argument.

## Implementing functional enrichment methods

Given a datsaset, we removed all genes from the annotation file that were not included in the dataset. Next, we deleted categories that annotated fewer than 20 or more than 500 genes included in the dataset. Categories that are too large are not informative, whereas categories that are too small have a substantial chance to be randomly enriched. Whenever such an upper and/or lower cutoff on the categories size is used, the exact composition of the investigated set of categories thus depends on the dataset. Removal of these categories reduces the search space, leading to faster convergence of the MCMC process.

Next, we deleted any genes that were not annotated to any categories (typically, none are deleted in this step) and sorted the list of remaining genes in descending order based on their differential expression levels (*log*2-fold change in the four datasets). This ranked gene list constitutes the main input to the functional enrichment method besides the annotation file. The dataset-specific size-filtering for categories is automatically implemented in CRFE and MGSA, but not for GSEA and ORA; therefore, we manually performed the filtering before running the latter two methods.

We used the CRFE software to run MGSA (with belief *b* = 1) and CRFE (with belief *b >* 1) and the R package `clusterProfiler` for GSEA and ORA [19]. Specifically, GSEA and ORA were performed using `clusterProfiler::GSEA` with fGSEA option [26] and `clusterProfiler::enricher`, which enable users to provide a custom annotation file. Because ORA is deterministic, we only ran it once for each dataset.

While GSEA includes a stochastic permutation test to compute the *p*-values, its main calculation of the Enrichment Scores is deterministic. Repeated GSEA runs showed minimal differences in the order of enriched categories and *p*-values; therefore, we only ran GSEA once for each dataset. In contrast, the MCMC process underlying MGSA and CRFE is inherently stochastic, we therefore repeated these analyses 100 times.

Except for GSEA, all methods require a threshold to divide the gene list into perturbed and unperturbed genes. Rather than picking a specific differential expression threshold *τ*, we chose *τ* such that, for each dataset, 30% of all genes were perturbed, and used 10% and 20% in sensitivity analyses. Depending on the dataset, the default threshold of 30% corresponds to an expression fold change threshold *τ* of 1.44-1.62, which is close to the frequently used value of 1.5 [37, 38] (Table 1).

**Table 1.** Gene list proportion and corresponding fold change. For the four datasets (rows) and different proportions of genes that are considered perturbed (columns), the table displays the minimal fold change values between treatment and control samples of the perturbed genes.

|  | 0.1 | 0.2 | 0.3 | 0.4 | 0.5 | 0.6 | 0.7 | 0.8 | 0.9 |
| --- | --- | --- | --- | --- | --- | --- | --- | --- | --- |
| **GSE40419** | 2.45 | 1.71 | 1.45 | 1.32 | 1.23 | 1.16 | 1.08 | 1.02 | 0.96 |
| **GSE87340** | 2.36 | 1.68 | 1.44 | 1.31 | 1.21 | 1.14 | 1.08 | 1.02 | 0.97 |
| **EENADCL** | 2.14 | 1.74 | 1.52 | 1.32 | 1.15 | 1.07 | 0.93 | 0.87 | 0.76 |
| **EENADLU** | 2.46 | 1.87 | 1.62 | 1.41 | 1.23 | 1.15 | 1.07 | 1.00 | 0.87 |

In terms of result interpretation, MGSA and CRFE return the enriched categories ranked by average posterior probability, GSEA ranks the enriched categories by normalized enrichment score (NES) or *p*-value, and ORA’s ranking is based on *p*-value. Therefore, we compared the results of seven methods: ORA, GSEA (ranked by NES and by *p*-value), MGSA, and CRFE with belief parameters 2, 5, and 10. We kept all categories enriched by GSEA and ORA by not providing any *p*-value cutoff on the results in order to capture all information embedded in the list of returned categories for the quality plots (see below). Unlike MGSA and CRFE which find enriched categories that explain many over-expressed genes, GSEA looks for categories in both directions, that is, categories that explain many over-expressed genes (positive NES) or many under-expressed genes (negative NES). For a fair comparison, we only considered GSEA-enriched categories with positive NES. We also filtered all categories with negative NES from the list of enriched categories ranked by GSEA *p*-value.

In addition, we post-processed the results of GSEA and ORA using REVIGO, as these two methods are prone to return many overlapping categories [28]. We used the REVIGO web interface with Homo sapiens Gene Ontology option at a medium allowed similarity (default) and SimRel as semantic similarity measure (default), and obsolete categories were not replaced. REVIGO-processed results contained fewer categories as each group of overlapping categories was collapsed into one representative category; we sorted the collapsed categories using the same ranking (*p*-value or NES) as the original list.

### Metrics for benchmarking functional enrichment methods

#### Redundancy

A common problem of functional enrichment methods that treat functional categories individually is a large overlap among the enriched categories in the explanatory set, i.e., multiple returned categories share many gene annotations, making the interpretation difficult. To quantify the redundancy of a set *C* of categories (returned by an algorithm), we define the *average redundancy R*(*C*) as the mean pairwise Jaccard similarity. That is,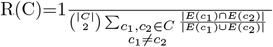 where *E* describes the set of all genes annotated to a given category.

#### Category size

Generally, functional categories with fewer gene annotations are more informative [15]. Therefore, a good method should preferably return specific categories. The biological processes included in this study were limited to having 20 to 500 gene annotations, and averaged approximately 100 gene annotations. For each method, we examined the number of genes annotated to the top returned categories, which we also call *category size*.

#### Quality measures

Functional enrichment methods seek to find a set of functional categories that annotate as many perturbed genes and as few unperturbed genes as possible. These two goals conflict with each: an improvement in one usually leads to a deterioration in the other. To quantify this trade-off, we defined a new measure, which we call quality. Given a set *C* of categories and a subset *X* of the perturbed genes *P*, the *quality Q*(*C, X*) of *C* with respect to *X* is the ratio of (a) the proportion of genes in *X* that are explained by *C* and (b) the proportion of unperturbed genes that are explained by *C*. That is,

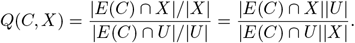

By defining the quality for any subset of the perturbed genes *P*, we can ask how it varies for different prefixes of the perturbed gene list (i.e., the top *x*% of *P*). If *C* = *C*, the quality is 1 as all genes are explained. If *C* does not explain any unperturbed genes, the quality equals infinity although for |*C*| *≥* 20 and with |*U* | = 70% of all genes, this never occurred in our study.

Given a ranked list of categories returned by a functional enrichment method, we created a diagnostic plot, the *quality plot*, as follows. Starting with the highest-ranked category as *C*, we plotted the quality *Q*(*C, X*) against the proportion of genes in *X* that are explained by *C*, |*E*(*C*) *∩ X*|*/*|*X*|, and repeated this as we added categories to *C* one at a time. We extrapolated each quality curve to the left boundary (i.e., 0 genes explained) with the same quality score as the first returned category and linearly interpolated to the right boundary at (1, 1), which is achieved when *C* = *C*. We used the area under the curve (AUC), computed with the trapezoidal rule, to compare the quality across different methods for different subsets *X* and for different proportions of perturbed genes.

Contrary to standard diagnostic plots such as the receiver operating characteristic or the precision-recall plot, a subset of perturbed genes *P − X* is not taken into account in the calculation of *Q*(*C, X*), unless *X* = *P*. This is so that the quality, when computed for different *X*, can highlight how well a method explains the highly perturbed genes without affecting the quality scores if these methods capture less perturbed genes.

## Results

Using the biological process aspect of the Gene Ontology and four published RNA-seq treatment-control differential gene expression datasets, we computed three metrics to evaluate the performance of CRFE and three established functional enrichment methods (MGSA, GSEA, and ORA) in (i) returning a small set of non-overlapping functional categories (conciseness), (ii) returning specific categories with few annotations (category size) and (iii) focusing on explaining highly perturbed genes (quality).

MGSA is a model-based approach that considers the enrichment of sets of functional categories but does not consider the ranking of the genes. CRFE generalizes MGSA by using information available in the ranking. GSEA uses the gene ranking as well but evaluates the enrichment of categories on an individual basis. ORA does not use the ranking of genes, and as GSEA, ORA considers each category individually. GSEA and ORA represent the two most frequently used methods. REVIGO is applied as a post-processing step to results from GSEA and ORA to find a representative, less redundant sub-list of enriched categories.

For CRFE, we tested three values for the belief parameter *b* = 2, 5, and 10, where a larger value puts more emphasis on highly perturbed genes. For all threshold-based methods (ORA, MGSA, and CRFE), we considered 30% of all genes perturbed (corresponding to perturbed genes exhibiting a fold-change of 1.44 or higher, Table 1). For GSEA, we ranked the enriched biological processes both by *p*-value (as frequently done by users) and by their normalized enrichment score (NES, which we show is superior). Lastly, we only considered GO biological processes that annotate between 20 *−* 500 genes included in the expression data.

## Conciseness

While GSEA and ORA assign similar enrichment values (*p*-value or NES) to categories with highly similar gene annotations, both MGSA and CRFE are, by design, more selective. Due to penalization for the number of enriched categories in the maximum likelihood function, MGSA and CRFE only add a category to the explanatory set if the added category annotates several perturbed genes not yet explained by the current explanatory set. Which one of several similar categories is ultimately preferred can depend on nuances such as the specific annotations and the categories already included in the explanatory set. This is also the reason why CRFE, in a preprocessing step, combines categories with the exact same gene annotations into one category. For three out of the four datasets, both GSEA and ORA (when using an unadjusted *p*-value cutoff of 0.05) returned more than twice the number of categories than MGSA or CRFE (when considering any category with an average posterior probability of 1% or higher; Figure 3). Even after a Benjamini-Hochberg correction for multiple testing, GSEA and ORA were on average less selective than CRFE.

**Fig 3.**
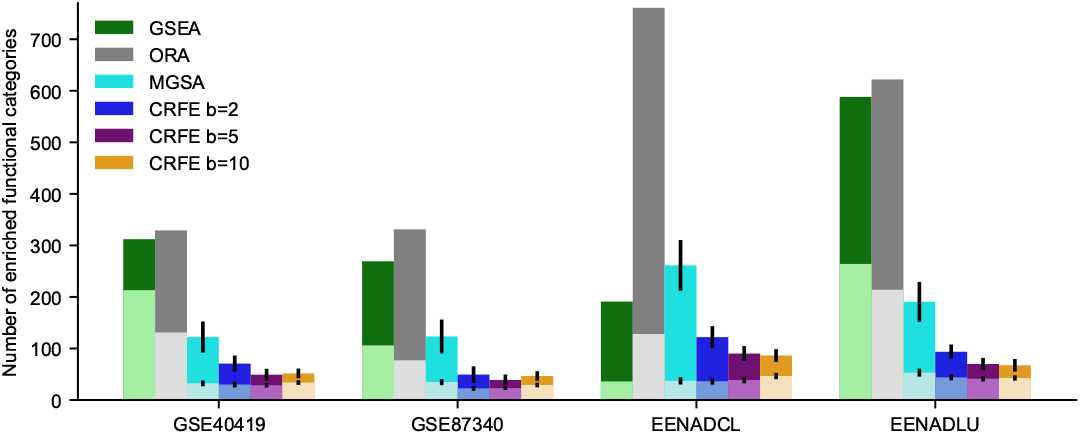
Number of returned functional categories. For each method and the four datasets (x-axis), the grouped bar chart shows the number of enriched functional categories. For GSEA and ORA, overlapping darker and lighter bars show the number of categories with *p*-value lower than 0.05 (before and after default Benjamini-Hochberg multiple testing adjustment). For MGSA and CRFE, overlapping darker and lighter bars show the number of categories with an average posterior higher than 1% and 10%, respectively. Error bars depict the standard deviations across 100 runs of MGSA and CRFE.

Among the two model-based methods, CRFE appears even more selective. Across the four datasets, MGSA consistently returned the most categories with an average posterior higher than 1%, followed by CRFE *b* = 2; CRFE *b* = 5 and *b* = 10, which returned roughly the same and lowest number of categories (Figure 3). This is likely due to the specifics of the MCMC process. To avoid getting trapped in a local optimum, the MCMC process accepts any proposal (e.g., the addition of any category) with a positive probability. However, the acceptance probability (Eq 3) decreases as the belief parameter increases, rendering the MCMC process more restrictive. Interestingly, the 13/27 number of categories with an average posterior higher than 10% (lighter bars of MGSA and CRFE in Figure 3) did not vary substantially when changing the belief parameter from 1 (MGSA) to 10. Categories with an average posterior of 10% or higher are likely truly enriched for perturbed genes, as a randomly added category is typically long removed before it accumulates 10% average posterior. Differences in the acceptance probability due to the belief parameter are small enough to barely affect categories with a true signal.

Researchers analyzing functional enrichment results often face long lists of functional categories that possess highly similar gene annotations due to the hierarchical structure of annotation databases such as the GO. These redundant lists are difficult to interpret and inflate the number of biologically meaningful results [28]. The average redundancy of a set of categories is measured by the average Jaccard similarity of gene annotations between all pairs of categories within the set. The higher the mean pairwise Jaccard similarity, the higher the average redundancy, and the more redundant a set of categories is. For all four datasets, the set of all biological processes, which annotate between 20 and 500 genes included in the dataset, is highly similar. The overall average redundancy of this entire set (i.e., the expected redundancy when randomly picking two categories) is 0.007-0.008 across all datasets.

For a fair comparison, we computed the average redundancy of the top 20 categories returned by each method. As expected, post-processing of GSEA and ORA results by REVIGO removed some highly redundant categories from the list and thus generally lowered the average redundancy (Figure 4). Only the GSEA NES results on the EENADCL dataset could not be improved by REVIGO as they already featured hardly any gene overlap between the top 20 returned categories before REVIGO application. REVIGO-improved GSEA and ORA enriched categories possessed, however, still more similar gene annotations than those returned by MGSA and CRFE. The simultaneous consideration of sets of categories as well as the penalization for the length of the explanatory set of categories allows CRFE and MGSA to avoid returning similar processes, to the point that their redundancy was as or almost as low as the expected redundancy. On the contrary, even a post-hoc REVIGO correction failed to achieve that level of conciseness.

**Fig 4.**
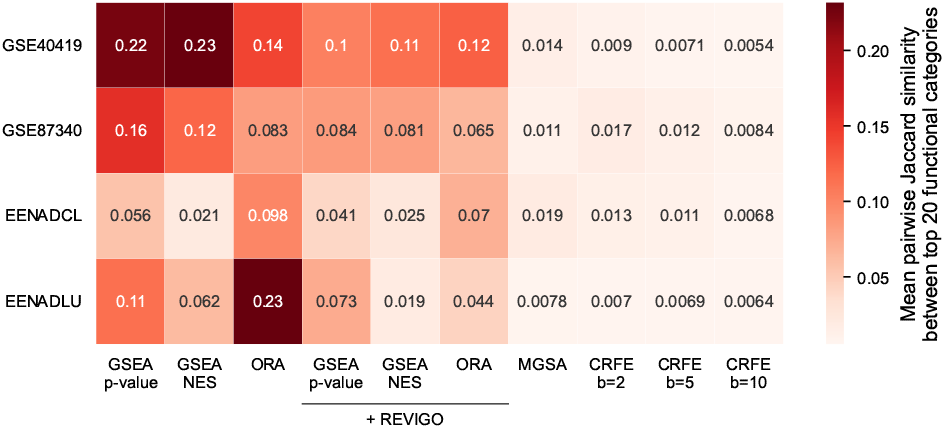
Average redundancy. For each method (x-axis) and each dataset (y-axis), the heatmap shows the mean pairwise Jaccard similarity in gene annotations of the top 20 returned categories, with more redundant returned sets of categories being shaded in a darker red. For all threshold-based methods, 30% of genes were considered perturbed.

## Category size

A good functional enrichment method preferably returns specific categories, i.e., categories with few gene annotations. These categories represent biological phenomena that are more clearly defined and thus more informative. To compare the specificity of the different methods, we examined how many measured genes were annotated to the respective top 20 returned biological processes. Across the four datasets, CRFE - irrespective of the chosen belief parameter - consistently returned smaller categories than other methods (Figure 5). GSEA NES and MGSA performed slightly worse and returned biological processes with an average category size that was similar to the average size of all considered processes with 20 to 500 annotations included in the respective dataset. Methods that ranked categories by the *p*-value (GSEA and ORA) predominantly returned larger categories, likely due to the well-known fact that the *p*-value decreases when “sample size” increases, even when the “effect size” remains fixed [17, 39]. The EENADCL dataset formed an exception in that all methods except for ORA, even GSEA *p*-value, returned quite specific categories. Post-hoc processing by REVIGO did not substantially alter the number of genes annotated to the top returned categories. This makes sense since REVIGO does not consider category size when selecting a representative category for a set of highly similar categories.

**Fig 5.**
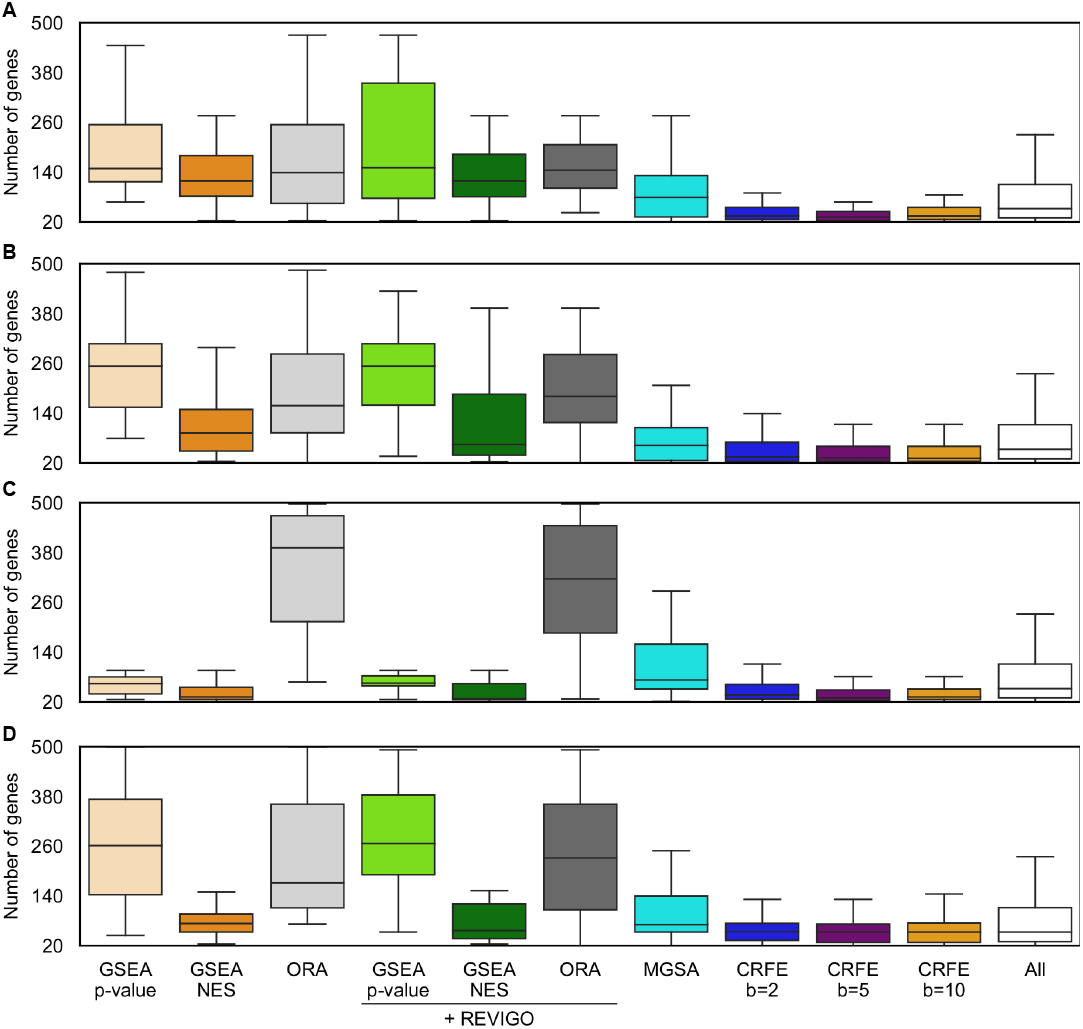
Size of the returned categories. For each method (x-axis) and each dataset (A: GSE40419, B: GSE87340, C: EENADCL, D: EENADLU), the boxplot shows the distribution of the number of genes annotated to the top 20 returned categories. For all threshold-based methods, 30% of genes were considered perturbed. Black lines depict the median, each box extends across the interquartile range (IQR), whiskers extend to the lowest data point still within 1.5 IQR of the lower quartile, and the highest data point still within 1.5 IQR of the upper quartile. Outliers are not displayed. The rightmost box depicts the distribution across all categories that possess 20 *−* 500 gene annotations included in the respective dataset.

## Quality

Functional enrichment methods should explain highly perturbed genes particularly well, as these genes are the most likely ones to be annotated to a functional category that was activated by the cell in response to some perturbation such as the development of cancer. Each method ranks the returned biological processes by some metric (e.g., *p*-value, average posterior). To compare how well each method explains the highest-ranked genes, we varied both the number *r* of categories included in the explanatory set and the percentage *x*% of genes considered highly ranked. As we varied *r*, we plotted the quality of the explanatory set against the proportion of top *x*% genes that was explained by the *r* categories. Figure 6A displays a representative quality plot for *x* = 15%. Recall that for the threshold-based methods, ORA, MGSA and CRFE, we considered 30% of all genes perturbed, corresponding to perturbed genes having a fold change of 1.45 or higher (Table 1). The choice *x* = 15% thus means that 50% of all perturbed genes were considered as positive in the quality computation. The remaining lower 50% perturbed genes are omitted from this computation. Quality scores before and after REVIGO application were very similar, which makes sense because REVIGO reduces a long list of categories by choosing representative categories and discarding semantically similar categories (which share many gene annotations). For visual reasons, we therefore only included REVIGO-corrected GSEA+REVIGO and ORA+REVIGO results in the quality plots.

**Fig 6.**
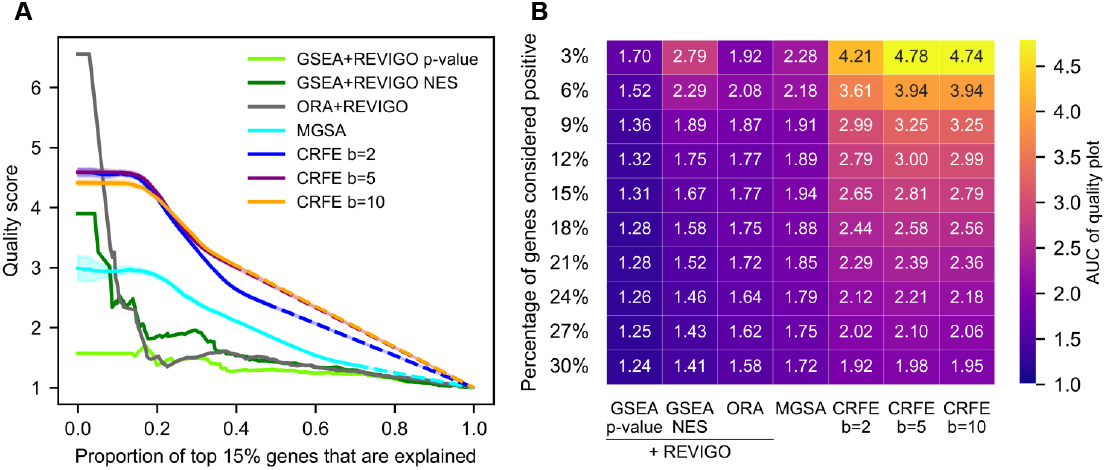
Quality of the returned sets of categories. For the GSE40419 dataset, the returned categories from each method are ranked based on NES score (GSEA), *p*-value (GSEA and ORA) or average posterior value (MGSA and CRFE). For all threshold-based methods, 30% of genes were considered perturbed. (A) For each method, categories are included one at a time and the quality in finding sets of categories that annotate top 15% genes versus bottom 70% genes (y-axis) is plotted against the proportion of top 15% genes that are annotated to an included category. Genes ranked at 15% *−* 30% were excluded from this computation. All stochastic methods (MGSA and CRFE) were repeated 100 times and shaded regions depict the 95%-confidence interval. For methods that do not return all categories (MGSA and CRFE), dotted lines depict a linear interpolation from the last computed point to the final value of (1, 1). (B) The area under the quality curve (as in A) is shown for different methods (x-axis) and for different proportions of genes that are considered positive (15% in A, y-axis).

Compared to MGSA, which does not take the ranking of perturbed genes into account, CRFE clearly succeeded in finding sets of categories that annotate more highly ranked genes at high quality (Figure 6A). The quality of ORA and GSEA, which consider each category individually, quickly dropped although these methods at least partially succeeded at finding initial categories with very high quality. For example, the first biological process returned by ORA+REVIGO is `DNA-templated DNA replication` with a quality of 6.6. This category is also selected by all other methods but not as the first one in the enriched list. Both MGSA and CRFE select this category within the first ten categories, with average posterior probabilities of 0.66 for MGSA and about 0.85 for CRFE, irrespective of the specific choice of tested belief parameter.

Next, we tested the sensitivity of these general trends by varying *x*, the percentage of genes considered positive, from 3% (i.e., 10% of all genes that are considered perturbed by ORA, MGSA, and CRFE) to 30% (i.e., 100%, respectively). To summarize the performance described by each quality plot, we computed the area under the curve (AUC). CRFE at *b* = 5 and *b* = 10 performed equally well (Figure 6B). However even at *x* = 30%, these two methods outperformed all other methods. For lower *x*, the difference in AUC values increased substantially, clearly highlighting that CRFE excels at explaining highly ranked or most perturbed genes. Furthermore, GSEA ranked by NES outperformed GSEA ranked by *p*-value. This is likely due to the preference for large categories when ranking categories by *p*-value, through which several undesired unperturbed genes are explained, lowering the quality. Notably, all methods explained the top ranked genes at highest quality, even ORA and MGSA, which do not consider the ranking of the perturbed genes. This provides evidence that one of the assumptions made by CRFE, namely that false positives appear more frequently at the lower end of the perturbed genes, is correct.

As a more comprehensive sensitivity analysis, we also varied the proportion of genes considered perturbed by threshold-based methods from 10% to 30%. Irrespective of the particular choice of this threshold, CRFE always returned sets of processes of highest quality (Figure 7), with the specific AUC values for the different methods and datasets tabulated in S1 Figure, S2 Figure, S3 Figure, and S4 Figure. This provides the most compelling argument for the superiority of CRFE over the other methods in focusing on explaining highly ranked genes. The higher CRFE’s belief parameter, the larger the emphasis CRFE places on explaining the top genes but, as a trade-off, the smaller the emphasis it places on explaining lowly perturbed genes. This explains why CRFE with *b* = 5 generally returned categories with higher quality than CRFE with *b* = 10 when focusing on explaining (i) most perturbed genes (e.g., all 100% perturbed genes) and/or (ii) many perturbed genes (e.g., 30%), while CRFE with *b* = 10 outperformed all other methods when focusing on the very top of the ranked gene list. The choice of the belief parameter should therefore ultimately depend on what proportion of genes a user wants to explain with high quality. As a rule of thumb, any value in the range of [5, 10] is a good choice, with the differences in AUC values between *b* = 5 and *b* = 10 being marginal.

**Fig 7.**
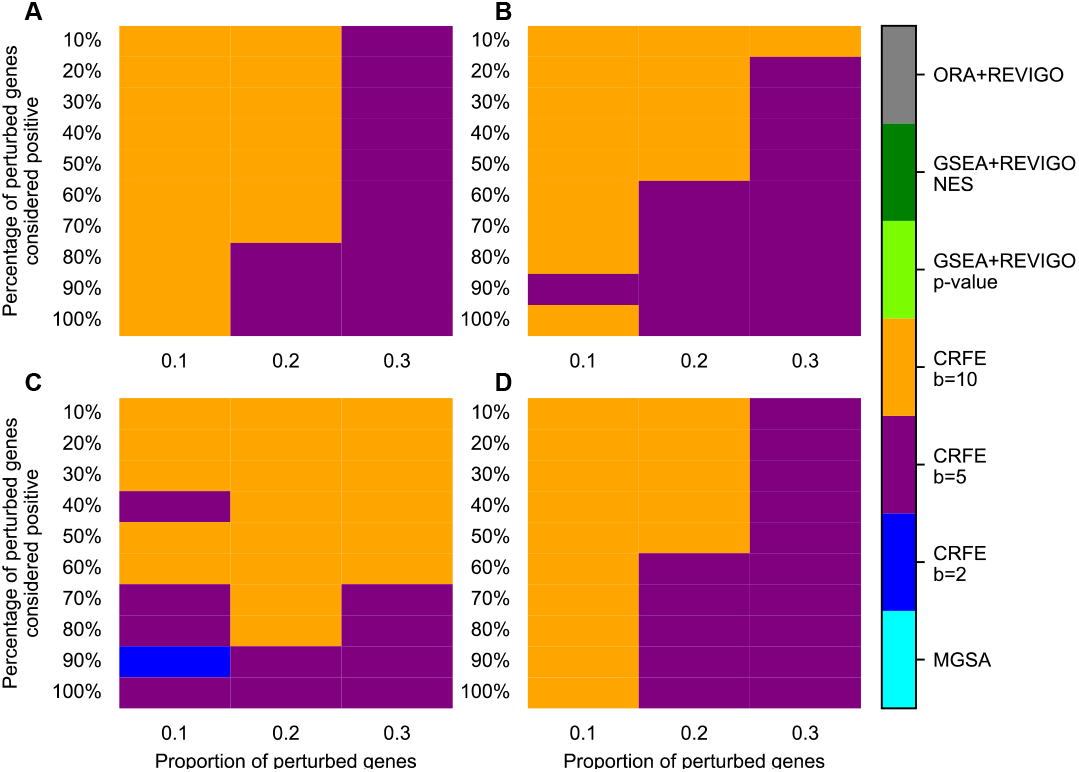
CRFE consistently returns sets of categories of highest quality. For various thresholds of genes that are considered perturbed (x-axis) as well as various proportions of perturbed genes used as positives in the quality plots (y-axis), the area under the quality curve (AUC, shown in Fig. 6A) is computed for all methods and the four datasets (A: GSE40419, B: GSE87340, C: EENADCL, D: EENADLU). Color indicates the method with the respectively highest AUC value. Exact AUC values for all methods are shown in S1 Figure - S4 Figure.

### Interpretation of enriched category sets

We performed a high-level interpretation of the results of CRFE for the two lung adenocarcinoma datasets. In our analysis we only considered positively differentially expressed genes as perturbed; therefore, each of the top 20 enriched biological processes reported in S1 Table and S2 Table represents activation in response to cancer. As expected, the set of the enriched processes for the two datasets is similar, indicating a consistent pattern of gene expression changes associated with lung adenocarcinoma. Most of the ten hallmarks of cancer appeared among the top enriched biological processes [40]. For instance, the enriched category `intermediate filament organization` alone is associated with eight hallmarks [41]. Categories such as `collagen metabolic process`, `alpha-amino acid biosynthetic process`, and `nucleoside diphosphate metabolic process` are relevant to the hallmarks of “Evading immune destruction” and “Activating invasion and metastasis” [42–44]. Additionally, “Chromosomal instability”, another hallmark, is represented by categories such as `centromere complex assembly`, `nuclear chromosome segregation`, and `DNA-templated DNA replication` [45–47]. Lastly, the hallmark “Deregulated metabolism” describes how cancer cells modify their cellular metabolic processes to sustain unlimited replication. Enriched categories such as `uronic acid metabolic process & glucuronate metabolic process`, `AMP metabolic process`, and `collagen catabolic process` are associated with this hallmark [42, 48, 49].

## Discussion

GSEA and ORA-based approaches constitute the most commonly used methods [6]. These methods regard each functional category individually, necessitating a false discovery rate control step that often eliminates a significant portion of the results. Moreover, these methods often return multiple categories with very similar gene annotations. Another method, MGSA, embeds all functional categories in a Bayesian network and analyzes them collectively. However, it only uses a binary classification of genes, disregarding valuable information contained in the gene ranking. Our new method CRFE, which generalizes MGSA by incorporating the specific ranking of genes, overcomes all these limitations.

CRFE contains two important hyperparameters: the belief and the proportion/threshold of perturbed genes. A higher belief parameter leads to a higher quality score when focusing on the small subset of most perturbed genes (Figures 6, 7). When the focus is on a larger proportion of perturbed genes, a lower belief parameter should be used.

Furthermore, a higher belief parameter yielded fewer enriched functional categories because the conditions imposed on acceptance of a category in the MCMC process became more restrictive (Table 3). Therefore, the choice of the belief parameter should depend on what proportion of perturbed genes a user wants to explain with a high quality. Experimalists often set the second hyperparameter, the threshold for perturbed genes, at expression fold changes of 1.5 or 2. When given the choice, we generally encourage users to choose looser thresholds since genes with a mediocre fold change still contain valuable information. Rather than choosing a specific fold change threshold, users can also simply consider a fixed proportion of all genes perturbed, as we did here. In evaluating the performance of functional enrichment methods in this study, we made an interesting observation about GSEA. Ranking the enriched functional categories by *p*-value versus NES value yielded on average larger and more redundant categories, which explained the perturbed genes at a lower quality. This is likely due to the bias of the *p*-value towards larger categories due to their increased statistical power. On the other hand, the NES value is not biased towards larger categories.

In conclusion, we showed that CRFE consistently outperforms the other methods when considering three key criteria. While all other methods exhibited poor performance in at least one criterion, CRFE returns a concise and specific set of functional categories that effectively explains highly perturbed genes with high quality. Using the adenocarcinoma data, we highlighted that the biological processes enriched by CRFE are biologically meaningful. Further exploration by domain experts would likely uncover additional interesting findings.

## Author Contributions

Conceptualization & Software: C.K.; Formal Analysis & Methodology & Writing: X.J, A.P. and C.K.

## Thanks

The authors thank T.M. Murali (Virginia Tech) for introducing them to the topic of functional enrichment analysis and initial fruitful discussions.

## Data and materials availability

Data and all relevant code is available at GitHub: https://github.com/ckadelka/CRFE.

## Supporting information

**S1 Figure.**
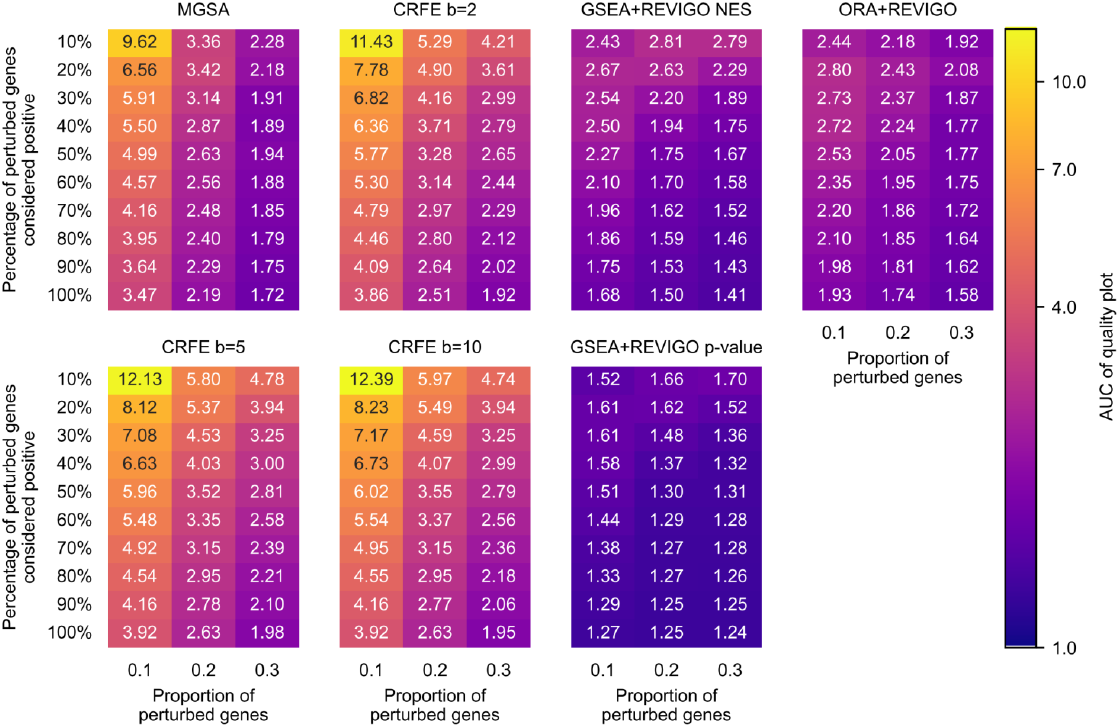
Area under the quality plot for the GSE40419 dataset.

**S2 Figure.**
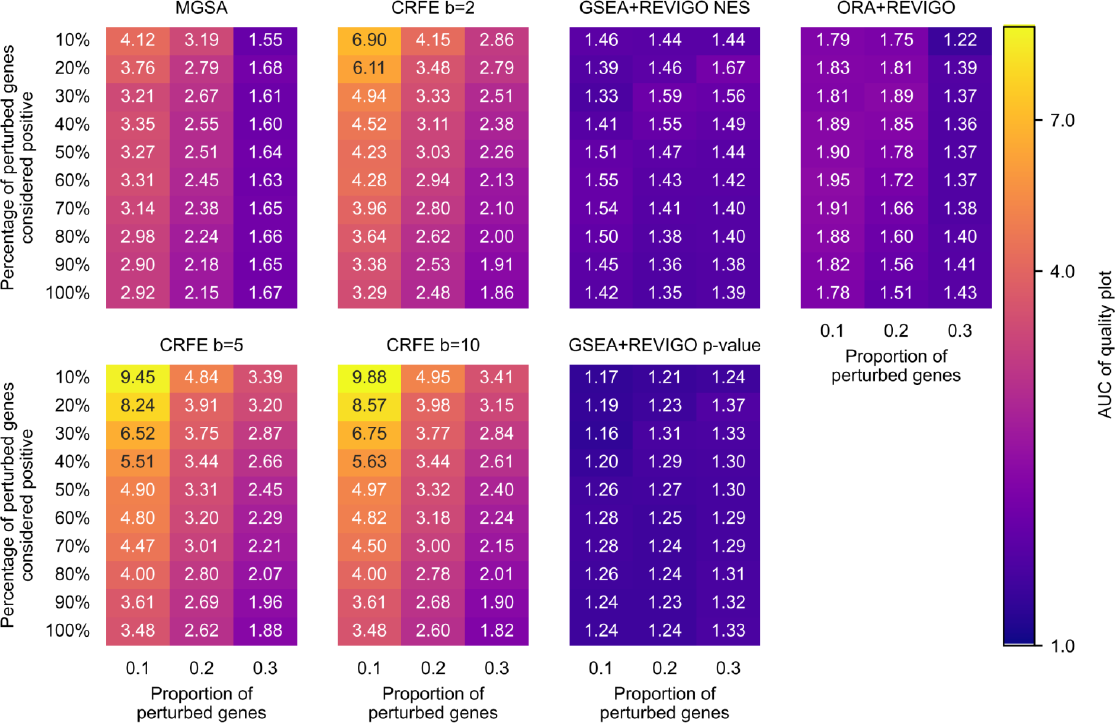
Area under the quality plot for the GSE87340 dataset.

**S3 Figure.**
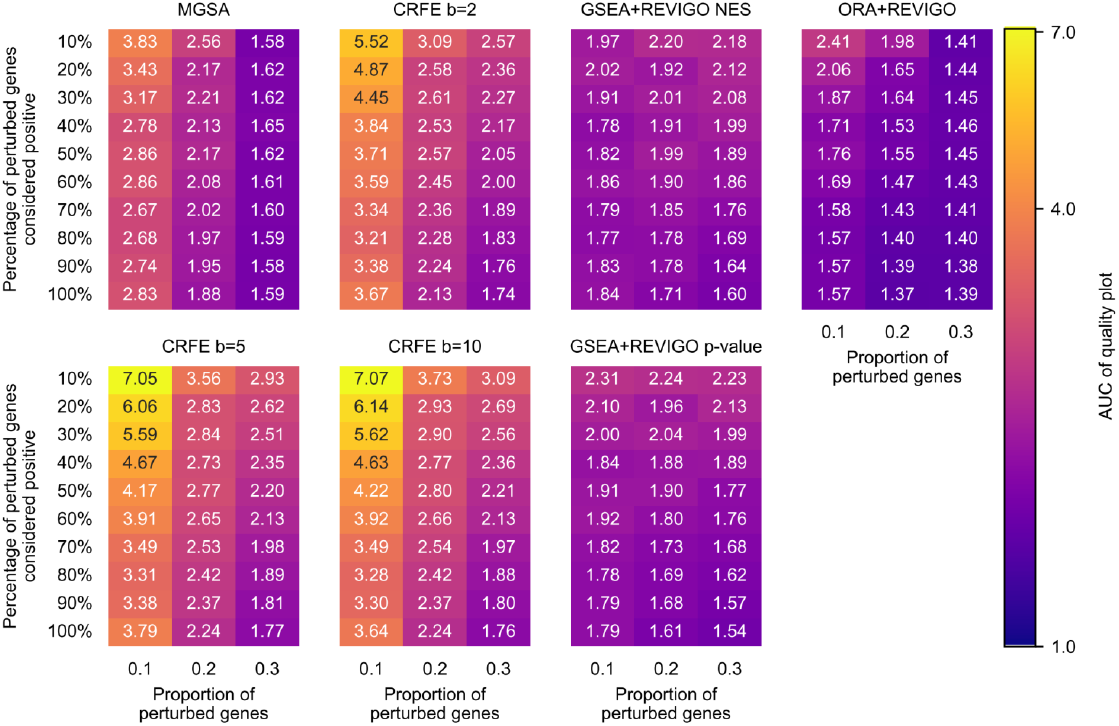
Area under the quality plot for the EENADCL dataset.

**S4 Figure.**
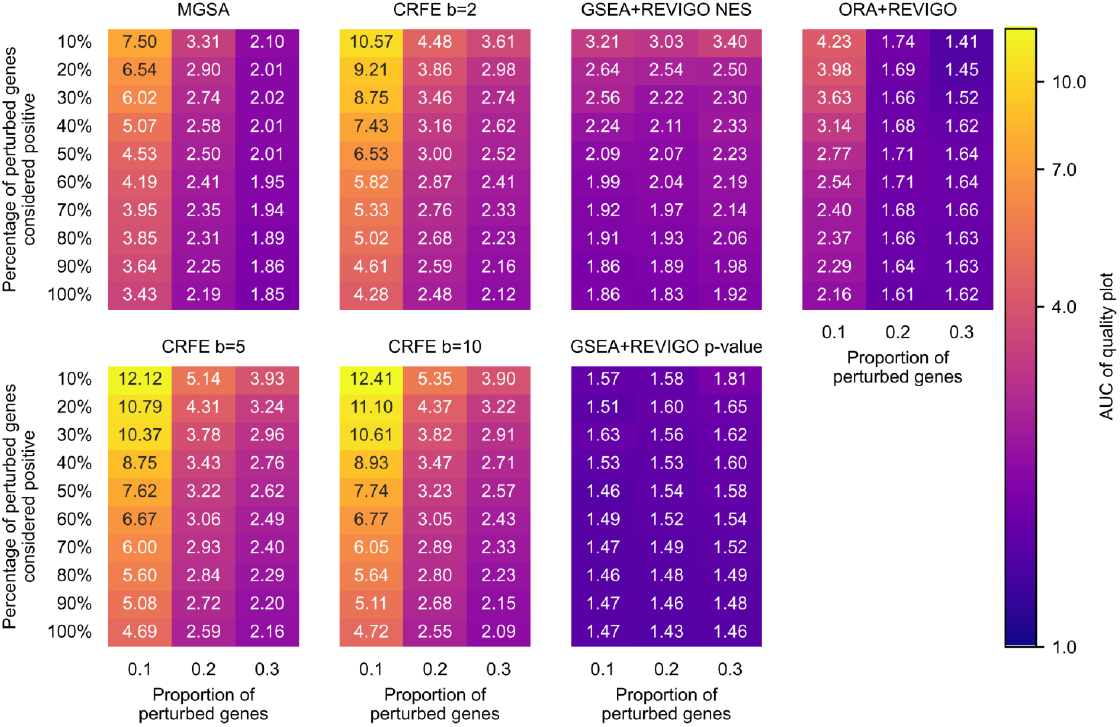
Area under the quality plot for the EENADLU dataset.

**S1 Table.** Top 20 biological processes returned by CRFE *b* = 5 for the GSE40419 dataset. 30% of all genes were considered perturbed and the MCMC process was run 100 times.

| rank | term | average posterior | stdev |
| --- | --- | --- | --- |
| 1 | digestion | 1.000 | 0 |
| 2 | uronic acid metabolic process & glucuronate metabolic process | 0.999 | 0.005 |
| 3 | centromere complex assembly | 0.998 | 0.024 |
| 4 | intermediate filament organization | 0.973 | 0.071 |
| 5 | antibacterial humoral response | 0.888 | 0.192 |
| 6 | nuclear chromosome segregation | 0.879 | 0.314 |
| 7 | sensory perception of taste | 0.864 | 0.192 |
| 8 | DNA-templated DNA replication | 0.832 | 0.344 |
| 9 | establishment of skin barrier | 0.774 | 0.279 |
| 10 | AMP metabolic process | 0.750 | 0.313 |
| 11 | xenobiotic transport | 0.730 | 0.262 |
| 12 | collagen catabolic process | 0.704 | 0.391 |
| 13 | nucleoside diphosphate metabolic process | 0.668 | 0.346 |
| 14 | neuron fate specification | 0.467 | 0.296 |
| 15 | regulation of kidney development | 0.466 | 0.299 |
| 16 | binding of sperm to zona pellucida | 0.451 | 0.314 |
| 17 | keratinization | 0.445 | 0.309 |
| 18 | positive regulation of G2/M transition of mitotic cell cycle | 0.341 | 0.309 |
| 19 | sperm-egg recognition | 0.318 | 0.275 |
| 20 | collagen metabolic process | 0.297 | 0.391 |

**S2 Table.** Top 20 biological processes returned by CRFE *b* = 5 for the GSE87340 dataset. 30% of all genes were considered perturbed and the MCMC process was run 100 times.

| rank | term | average posterior | stdev |
| --- | --- | --- | --- |
| 1 | AMP metabolic process | 0.995 | 0.034 |
| 2 | keratinization | 0.990 | 0.064 |
| 3 | intermediate filament organization | 0.987 | 0.067 |
| 4 | centromere complex assembly | 0.907 | 0.186 |
| 5 | establishment of skin barrier | 0.873 | 0.227 |
| 6 | digestion | 0.861 | 0.307 |
| 7 | uronic acid metabolic process & glucuronate metabolic process | 0.841 | 0.321 |
| 8 | alpha-amino acid biosynthetic process | 0.692 | 0.315 |
| 9 | regulation of branching involved in ureteric bud morphogenesis | 0.590 | 0.377 |
| 10 | collagen metabolic process | 0.550 | 0.414 |
| 11 | cell cycle DNA replication | 0.495 | 0.389 |
| 12 | nuclear DNA replication | 0.480 | 0.384 |
| 13 | negative regulation of nuclear division | 0.461 | 0.418 |
| 14 | regulation of mitotic nuclear division | 0.451 | 0.417 |
| 15 | chromosome condensation | 0.414 | 0.297 |
| 16 | columnar/cuboidal epithelial cell differentiation | 0.399 | 0.357 |
| 17 | collagen catabolic process | 0.378 | 0.403 |
| 18 | response to dietary excess | 0.314 | 0.248 |
| 19 | metanephric nephron morphogenesis | 0.239 | 0.311 |
| 20 | neuroepithelial cell differentiation | 0.230 | 0.283 |

**S3 Table.** Top 20 biological processes returned by CRFE *b* = 5 for the EENADCL dataset. 30% of all genes were considered perturbed and the MCMC process was run 100 times.

| rank | term | average posterior | stdev |
| --- | --- | --- | --- |
| 1 | vasculogenesis | 0.994 | 0.023 |
| 2 | lymph vessel development | 0.902 | 0.188 |
| 3 | negative regulation of BMP signaling pathway | 0.875 | 0.313 |
| 4 | regulation of cytoplasmic transport | 0.781 | 0.234 |
| 5 | response to thyroid hormone | 0.781 | 0.233 |
| 6 | positive regulation of glucose transmembrane transport | 0.745 | 0.318 |
| 7 | synaptic vesicle transport | 0.567 | 0.26 |
| 8 | cyclic-nucleotide-mediated signaling | 0.550 | 0.406 |
| 9 | renal system vasculature development & kidney vasculature development | 0.460 | 0.316 |
| 10 | regulation of cytoplasmic translation | 0.453 | 0.271 |
| 11 | negative regulation of calcium ion transmembrane transporter activity | 0.434 | 0.315 |
| 12 | regulation of focal adhesion assembly & regulation of cell-substrate junction assembly | 0.424 | 0.403 |
| 13 | cGMP-mediated signaling | 0.422 | 0.405 |
| 14 | cellular response to hypoxia | 0.415 | 0.376 |
| 15 | chaperone-mediated protein complex assembly | 0.395 | 0.233 |
| 16 | maturation of LSU-rRNA | 0.385 | 0.203 |
| 17 | regulation of cell-substrate junction organization | 0.376 | 0.406 |
| 18 | glomerulus vasculature development | 0.353 | 0.266 |
| 19 | negative regulation of calcium ion transmembrane transport | 0.322 | 0.294 |
| 20 | mitochondrion morphogenesis | 0.301 | 0.254 |

**S4 Table.** Top 20 biological processes returned by CRFE *b* = 5 for the EENADLU dataset. 30% of all genes were considered perturbed and the MCMC process was run 100 times.

| rank | term | average posterior | stdev |
| --- | --- | --- | --- |
| 1 | collagen metabolic process | 0.956 | 0.178 |
| 2 | chondrocyte development | 0.940 | 0.165 |
| 3 | folic acid-containing compound metabolic process | 0.876 | 0.287 |
| 4 | chromosome condensation | 0.849 | 0.336 |
| 5 | antimicrobial humoral immune response mediated by antimicrobial peptide | 0.841 | 0.337 |
| 6 | response to vitamin | 0.813 | 0.323 |
| 7 | cytoplasmic translation | 0.770 | 0.369 |
| 8 | cellular oxidant detoxification | 0.729 | 0.403 |
| 9 | acute inflammatory response | 0.705 | 0.394 |
| 10 | ATP synthesis coupled electron transport & mitochondrial ATP synthesis coupled electron transport | 0.637 | 0.441 |
| 11 | response to copper ion | 0.627 | 0.323 |
| 12 | protein hydroxylation | 0.624 | 0.346 |
| 13 | negative regulation of endopeptidase activity | 0.591 | 0.323 |
| 14 | chaperone-mediated protein folding | 0.540 | 0.491 |
| 15 | positive regulation of macrophage activation | 0.510 | 0.342 |
| 16 | collagen fibril organization | 0.431 | 0.412 |
| 17 | positive regulation of DNA biosynthetic process | 0.405 | 0.481 |
| 18 | protein refolding | 0.374 | 0.404 |
| 19 | negative regulation of viral life cycle | 0.323 | 0.302 |
| 20 | proton motive force-driven ATP synthesis | 0.323 | 0.392 |

## References

1. Reuter JA, Spacek DV, Snyder MP. High-throughput sequencing technologies. Mol Cell. 2015;58(4):586–597.

2. Mardis ER. The impact of next-generation sequencing technology on genetics. Trends Genet. 2008;24(3):133–141.

3. Yadav SP. The wholeness in suffix-omics, -omes, and the word om. J Biomol Tech. 2007;18(5):277.

4. Sandhu C, Qureshi A, Emili A. Panomics for Precision Medicine. Trends Mol Med. 2018;24(1):85–101.

5. Wijesooriya K, Jadaan SA, Perera KL, Kaur T, Ziemann M. Urgent need for consistent standards in functional enrichment analysis. PLoS Comput Biol. 2022;18(3):e1009935.

6. Maleki F, Ovens K, Hogan DJ, Kusalik AJ. Gene Set Analysis: Challenges, Opportunities, and Future Research. Front Genet. 2020;11:654.

7. Garcia-Moreno A, Lòpez-Domínguez R, Villatoro-García JA, Ramirez-Mena A, Aparicio-Puerta E, Hackenberg M, et al. Functional Enrichment Analysis of Regulatory Elements. Biomedicines. 2022;10(3).

8. Chung M, Bruno VM, Rasko DA, Cuomo CA, Muñoz JF, Livny J, et al. Best practices on the differential expression analysis of multi-species RNA-seq. Genome Biol. 2021;22(1):121.

9. Love MI, Huber W, Anders S. Moderated estimation of fold change and dispersion for RNA-seq data with DESeq2. Genome Biol. 2014;15(12):550.

10. Wang K, Li M, Bucan M. Pathway-based approaches for analysis of genomewide association studies. Am J Hum Genet. 2007;81(6):1278–1283.

11. Pang Z, Chong J, Zhou G, de Lima Morais DA, Chang L, Barrette M, et al. MetaboAnalyst 5.0: narrowing the gap between raw spectra and functional insights. Nucleic Acids Res. 2021;49(W1):W388–W396.

12. Schölz C, Lyon D, Refsgaard JC, Jensen LJ, Choudhary C, Weinert BT. Avoiding abundance bias in the functional annotation of post-translationally modified proteins. Nat Methods. 2015;12(11):1003–1004.

13. Ashburner M, Ball CA, Blake JA, Botstein D, Butler H, Cherry JM, et al. Gene ontology: tool for the unification of biology. The Gene Ontology Consortium. Nat Genet. 2000;25(1):25–29.

14. Mazandu GK, Mulder NJ. Information content-based gene ontology semantic similarity approaches: toward a unified framework theory. BioMed research international. 2013;2013.

15. Resnik P. Semantic Similarity in a Taxonomy: An Information-Based Measure and its Application to Problems of Ambiguity in Natural Language. Journal of Artificial Intelligence Research. 1999;11:95–130.

16. Mazandu GK, Mulder NJ. Information content-based gene ontology semantic similarity approaches: toward a unified framework theory. Biomed Res Int. 2013;2013:292063.

17. Khatri P, Sirota M, Butte AJ. Ten Years of Pathway Analysis: Current Approaches and Outstanding Challenges. PLOS Computational Biology. 2012;8(2):e1002375. Publisher: Public Library of Science.

18. Subramanian A, Tamayo P, Mootha VK, Mukherjee S, Ebert BL, Gillette MA, et al. Gene set enrichment analysis: a knowledge-based approach for interpreting genome-wide expression profiles. Proceedings of the National Academy of Sciences of the United States of America. 2005;102(43):15545–15550.

19. Yu G, Wang LG, Han Y, He QY. clusterProfiler: an R package for comparing biological themes among gene clusters. Omics: a journal of integrative biology. 2012;16(5):284–287.

20. Reimand J, Kull M, Peterson H, Hansen J, Vilo J. g:Profiler – a web-based toolset for functional profiling of gene lists from large-scale experiments. Nucleic Acids Res. 2007;35(Web Server issue):193–200.

21. Fang Z, Liu X, Peltz G. GSEApy: a comprehensive package for performing gene set enrichment analysis in Python. Bioinformatics. 2023;39(1).

22. Sherman BT, Hao M, Qiu J, Jiao X, Baseler MW, Lane HC, et al. DAVID: a web server for functional enrichment analysis and functional annotation of gene lists (2021 update). Nucleic Acids Res. 2022;50(W1):W216–W221.

23. Efron B, Tibshirani R. On testing the significance of sets of genes. aoas. 2007;1(1):107–129.

24. Simillion C, Liechti R, Lischer HEL, Ioannidis V, Bruggmann R. Avoiding the pitfalls of gene set enrichment analysis with SetRank. BMC Bioinformatics. 2017;18(1):151.

25. Roder J, Linstid B, Oliveira C. Improving the power of gene set enrichment analyses. BMC Bioinformatics. 2019;20(1):257.

26. Korotkevich G, Sukhov V, Budin N, Shpak B, Artyomov MN, Sergushichev A. Fast gene set enrichment analysis. BioRxiv [Preprint]. 2021 [posted 2016 June 20; revised 2019 Oct 22; revised 2021 Feb 1; cited 2023 June 27];Available from: https://www.biorxiv.org/content/10.1101/060012v3.

27. Geistlinger L, Csaba G, Santarelli M, Ramos M, Schiffer L, Turaga N, et al. Toward a gold standard for benchmarking gene set enrichment analysis. Brief Bioinform. 2021;22(1):545–556.

28. Supek F, Bosnjak M, Skunca N, Smuc T. REVIGO summarizes and visualizes long lists of gene ontology terms. PloS one. 2011;6(7):e21800.

29. Fröhlich H, Speer N, Poustka A, Beissbarth T. GOSim–an R-package for computation of information theoretic GO similarities between terms and gene products. BMC bioinformatics. 2007;8:166.

30. Bauer S, Gagneur J, Robinson PN. GOing Bayesian: model-based gene set analysis of genome-scale data. Nucleic Acids Res. 2010;38(11):3523–3532.

31. Consortium GO. The Gene Ontology (GO) database and informatics resource. Nucleic acids research. 2004;32(suppl 1):D258–D261.

32. Seo JS, Ju YS, Lee WC, Shin JY, Lee JK, Bleazard T, et al. The transcriptional landscape and mutational profile of lung adenocarcinoma. Genome research. 2012;22(11):2109–2119.

33. Sun Z, Wang L, Eckloff BW, Deng B, Wang Y, Wampfler JA, et al. Conserved recurrent gene mutations correlate with pathway deregulation and clinical outcomes of lung adenocarcinoma in never-smokers. BMC Medical Genomics. 2014;7(1):1–11.

34. Wu M, Chen Y, Xia H, Wang C, Tan CY, Cai X, et al. Transcriptional and proteomic insights into the host response in fatal COVID-19 cases. Proceedings of the National Academy of Sciences of the United States of America. 2020;117(45):28336–28343.

35. Love MI, Huber W, Anders S. Moderated estimation of fold change and dispersion for RNA-seq data with DESeq2. Genome biology. 2014;15(12):1–21.

36. Papatheodorou I, Moreno P, Manning J, Fuentes AMP, George N, Fexova S, et al. Expression Atlas update: from tissues to single cells. Nucleic acids research. 2020;48(D1):D77–D83.

37. McCarthy DJ, Smyth GK. Testing significance relative to a fold-change threshold is a TREAT. Bioinformatics. 2009;25(6):765–771.

38. Dalman MR, Deeter A, Nimishakavi G, Duan ZH. Fold change and p-value cutoffs significantly alter microarray interpretations. BMC bioinformatics. 2012;13:1–4.

39. Sullivan GM, Feinn R. Using effect size—or why the P value is not enough. Journal of graduate medical education. 2012;4(3):279–282.

40. Hanahan D, Weinberg RA. Hallmarks of Cancer: The Next Generation. Cell. 2011;144(5):646–674.

41. Sharma P, Alsharif S, Fallatah A, Chung BM. Intermediate Filaments as Effectors of Cancer Development and Metastasis: A Focus on Keratins, Vimentin, and Nestin. Cells. 2019;8(5).

42. Xu S, Xu H, Wang W, Li S, Li H, Li T, et al. The role of collagen in cancer: from bench to bedside. J Transl Med. 2019;17(1):309.

43. Butler M, van der Meer LT, van Leeuwen FN. Amino Acid Depletion Therapies: Starving Cancer Cells to Death. Trends Endocrinol Metab. 2021;32(6):367–381.

44. Boison D, Yegutkin GG. Adenosine Metabolism: Emerging Concepts for Cancer Therapy. Cancer Cell. 2019;36(6):582–596.

45. Zhang CZ, Spektor A, Cornils H, Francis JM, Jackson EK, Liu S, et al. Chromothripsis from DNA damage in micronuclei. Nature. 2015;522(7555):179–184.

46. Wang X, Chen D, Gao J, Long H, Zha H, Zhang A, et al. Centromere protein U expression promotes non-small-cell lung cancer cell proliferation through FOXM1 and predicts poor survival. Cancer Manag Res. 2018;10:6971–6984.

47. Chow SE, Meir YJJ, Li JM, Hsu PC, Yang CT. Nuclear p120 catenin is a component of the perichromosomal layer and coordinates sister chromatid segregation during mitosis in lung cancer cells. Cell Death Dis. 2022;13(6):526.

48. Fahrmann JF, Grapov DD, Wanichthanarak K, DeFelice BC, Salemi MR, Rom WN, et al. Integrated Metabolomics and Proteomics Highlight Altered Nicotinamide- and Polyamine Pathways in Lung Adenocarcinoma. Carcinogenesis. 2017;38(3):271–280.

49. Moldogazieva NT, Mokhosoev IM, Terentiev AA. Metabolic Heterogeneity of Cancer Cells: An Interplay between HIF-1, GLUTs, and AMPK. Cancers. 2020;12(4).

